# Mitochondrial Genome Sequence of *Phytophthora sansomeana* and Comparative Analysis of *Phytophthora* Mitochondrial Genomes

**DOI:** 10.1101/2020.03.23.003319

**Authors:** Guohong Cai, Steven R. Scofield

## Abstract

*Phytophthora sansomeana* infects soybean and causes root rot. It was recently separated from the species complex *P. megasperma sensu lato.* In this study, we sequenced and annotated its complete mitochondrial genome and compared it to that of nine other *Phytophthora* species. The genome was assembled into a circular molecule of 39,618 bp with a 22.03% G+C content. Forty-two protein coding genes, 25 tRNA genes and two rRNA genes were annotated in this genome. The protein coding genes include 14 genes in the respiratory complexes, four ATP synthetase genes, 16 ribosomal proteins genes, a *tatC* translocase gene, six conserved ORFs and a unique *orf402.* The tRNA genes encode tRNAs for 19 amino acids. Comparison among mitochondrial genomes of 10 *Phytophthora* species revealed three inversions, each covering multiple genes. These genomes were conserved in gene content with few exceptions. A 3’ truncated *atp9* gene was found in *P. nicotianae.* All 10 *Phytophthora* species, as well as other oomycetes and stramenopiles, lacked tRNA genes for threonine in their mitochondria. Phylogenomic analysis using the mitochondrial genomes supported or enhanced previous findings of the phylogeny of *Phytophthora* spp.

## Introduction

The genus *Phytophthora* includes many devastating pathogens infecting economically important crops [1]. Perhaps the best known and most historically significant among those is *P. infestans,* the pathogen that causes potato late blight and the culprit behind the Irish potato famine [2]. During the past 20 years, the number of species described in the *Phytophthora* genus has expanded significantly. With approximately 55 species described in 1999, it expanded to 105 species by 2007 [3] and 117 species by 2012 [4]. Currently, 123 formally described species and 23 provisional species are included in the *Phytophthora* Database (www.phytophthoradb.org) [5]. *Phytophthora* is a genus in the *Peronosporales* of oomycetes. Oomycetes produce hyphae and are morphologically similar to fungi. However, they are phylogenetically distant. Oomycetes belong to the major group Stramenopila, which also include diatoms and brown algae [6–8].

### *P. sansomeana* is one of the recently described species in the *Phytophthora* genus

Previously included in the species complex *P. megasperma sensu lato,* it is morphologically similar to but phylogenetically distinct from *P. megasperma sensu strictu [9]. P. sansomeana* was isolated from soybean in Indiana in 1990 [10] and has now been reported in China [11], Canada and multiple Midwest states in the USA [12, 13]. It causes discoloration and rotting of lateral root and internal discoloration and rotting of the taproot. Above ground, it can cause yellowing and stunting, or whole plant wilting, but the chocolate-colored stem discoloration which is typically associated with late-season infection by another soybean root rot pathogen, *P. sojae,* is usually absent in *P. sansomeana-infected* soybean plants. Also included in this species are isolates from Douglas-fir and alfalfa [9].

Interest in the genus *Phytophthora* has been increasing due to its economic impact. Molecular data serves as the foundation of many studies. Genome sequences have been available for many *Phytophthora* species. Complete mitochondrial genomes from eight species have been published. These include *P. infestans* [14, 15], *P. andina, P. ipomoeae, P. mirabilis, P. phaseoli* [16], *P. nicotianae* [17], *P. sojae* and *P. ramorum* [18]. Another species, *P. polonica,* also has its complete mitochondrial genome publicly available in GenBank (Table 1). Inconsistencies in annotation have been observed. Mitochondrial genomes have been use to study the population of *P. infestans* [19–22] as well as resolving the phylogenetic relationship of *Phytophthora* species [15, 23]. Molecular information from *P. sansomeana* is limited. In this study, we sequenced and annotated its complete mitochondrial genome, compared it to that of other *Phytophthora* species currently publicly available, and examined phylogenetic relationship among these *Phytophthora* species using their mitochondrial genomes.

**Table 1.** Mitochondrial genomes of 10 *Phytophthora* species included in the analysis in this study

| Species | Accession # | Length (bp) | GC (%) | Clade<sup>a</sup> | Reference |
| --- | --- | --- | --- | --- | --- |
| *P. infestans* |  |  |  |  |  |
| Haplotype Ia | AY894835 | 37,922 | 22.32 | 1c | [15] |
| Ib | U17009 | 37,957 | 22.29 | 1c | [14] |
| IIa | AY898627 | 39,870 | 22.29 | 1c | [15] |
| IIb | AY898628 | 39,840 | 22.30 | 1c | [15] |
| *P. andina* |  |  |  |  |  |
| Haplotype Ia | KJ408269<sup>b</sup> | 37,883 | 22.29 | 1c | [16] |
| Ic | HM590419 | 37,874 | 22.14 | 1c | [16] |
| *P. ipomoeae* | HM590420 | 37,872 | 22.39 | 1c | [16] |
| *P. mirabilis* | HM590421 | 37,779 | 22.38 | 1c | [16] |
| *P. phaseoli* | HM590418 | 37,914 | 22.14 | 1c | [16] |
| *P. nicotianae* | KY851301 | 37,561 | 21.88 | 1 | [17] |
| *P. sojae* | DQ832717 | 42,977 | 21.70 | 7 | [18] |
| *P. polonica* | KT946598 | 40,467 | 21.57 | 9 | N/A |
| *P. ramorum* | DQ832718 | 39,314 | 21.98 | 8c | [18] |
| *P. sansomeana* | MH936679 | 39,618 | 22.03 | 8a | This study |

<sup>a</sup> The designation of phylogenetic clade is based on [23, 37].
<sup>b</sup> This genome is incomplete.

## Materials and Methods

### Nucleic acids extraction and sequencing

The type strain of *P. sansomeana,* 1819b, was obtained from American Type Culture Collection (ATCC #: MYA-4455). It was maintained on lima bean agar. For nucleic acids extraction, it was grown in half-strength lima bean broth on a lab bench for 8 days. DNA was extracted from harvested mycelium using the Gentra Puregene Yeast/Bact kit (Qiagen).

DNA was sequenced using both PacBio and Illumina technologies. For PacBio sequencing, a 20-kb insert library was constructed and then run on two SMRT cells on the RS II sequencing platform using the P6-C4 chemistry. For Illumina sequencing, a TruSeq DNA PCR-free library with mean insert size of approximately 440 bp was constructed according to manufacturer’s instructions. It was sequenced on the HighSeq2500 platform.

### Assembly, annotation and comparative genomics

The PacBio reads were assembled using HGAP (RS_HGAP_Assembly.3 protocol implemented in PacBio smrtanalysis 2.3.0 software package) and polished using Quiver [24]. Illumina read pairs were mapped onto the assembly using Bowtie2 [25], and Pilon [26] was used to further improve the assembly by utilizing the aligned Illumina reads.

The mitochondrial genome of *P. sansomeana* was first annotated using MFannot (http://megasun.bch.umontreal.ca/cgi-bin/mfannot/mfannotInterface.pl) using the standard genetic code. Like plant mitochondria, but different from those of animal and fungi, oomycete mitochondria use the standard code [27]. The automatic annotation was then verified manually. Mitochondrial genomes of nine other *Phytophthora* species were downloaded from GenBank (Table 1). Protein coding genes among the species were compared. Genes identified in one species but not in another were manually verified or corrected. For this purpose, ORFfinder (https://www.ncbi.nlm.nih.gov/orffinder/) was used to find open reading frames (ORFs) and BLAST searches were used to identify similarity. ORFfinder was also used to find ORFs encoding at least 100 aa in the intergenic regions of all *Phytophthora* mitochondrial genomes. Ribosomal RNA (rRNA) gene sequences were used to search NCBI non-redundant database and SILVA rRNA database [28]. Transfer RNA (tRNA) genes were analyzed using tRNAscan-SE 2.0 [29] in the organellar mode. A circular map of the mitochondrial genome of *P. sansomeana* was generated using OGDRAW [30]. It’s genome assembly and annotation were submitted to GenBank under accession number MH936679.

### Phylogenomic analysis

To construct phylogenetic tree based on the mitochondrial genomes, individual protein coding genes and rRNA genes were aligned with Muscle [31]. The alignments were visually inspected to remove gaps and regions deemed ambiguously aligned (usually between closely adjacent gaps). The alignments were concatenated and a maximum-likelihood tree based on the Tamura-Nei nucleotide model [32] was constructed using MEGA7 [33]. One thousand bootstraps were used to test the support of individual branches. *Pythium ultimum* [34] and *Pythium insidiosum* [35] were used as outgroup.

## Results and discussion

### Assembly and annotation of *P. sansomeana* mitochondrial genome

PacBio sequencing generated 383,775 subreads totaling 2.57 Gb after filtering and adapter removal. The N50 of subread length was 9,846 bp. The mitochondrial genome of *P. sansomeana* was assembled into a single contig of 55.5 kb using PacBio reads, with a 15.8 kb direct duplicated region at both ends, indicating a circular topology. To verify this, the contig was broken at a random position in the non-duplicated region and the two segments were reconnected by merging the duplicated regions. PacBio reads were then mapped to this rearranged contig. Visual inspection of read alignments and coverage confirmed its circular topology. After polishing, the PacBio assembly resulted in a molecule of 39,546 bp. Illumina sequencing generated 43.3 million read pairs at 151 bp x 2 in length. A total of 2,249,965 read pairs were mapped to the mitochondrial genome. Based on the alignments of Illumina reads, 72 single nucleotide deletions were corrected.

The mitochondrial genome of *P. sansomeana* was assembled into a circular molecule of 39,618 bp with a 22.03% G+C content. It was predicted to encode 42 protein coding genes, two ribosomal RNA genes, and 25 tRNA genes. The genes were encoded by both strands. The protein coding genes included 10 NADH dehydrogenase genes in respiratory complex I *(nad1, nad2, nad3, nad4, nad4L, nad5, nad6, nad7, nad9* and *nad11),* one cytochrome C reductase gene (cob) in complex III, and three cytochrome c oxidase genes in complex IV (cox1, *cox2* and *cox3).* Also included were four ATP synthase genes *(atp1, atp6, atp8* and *atp9),* five large subunit ribosomal protein genes *(rpl2, rpl5, rpl6, rpl14* and *rpl16),* and 11 small subunit ribosomal protein genes *(rps2, rps3, rps4, rps7, rps8, rps10, rps11, rps12, rps13, rps14* and *rps19)* (Fig. 1). We annotated a sec-independent translocase gene (tatC) in the twin-arginine translocation system. The annotation of this gene was inconsistent in other *Phytophthora* mitochondrial genomes (see analysis below). The six conserved ORFs with no annotated function found in other *Phytophthora* mitochondrial genomes (Table 1) were also found in *P. sansomeana (orf32, ymf96 (=orf79), ymf98 (=orf142), ymf99(=orf217), ymf100 (=orf100),* and *ymf101 (=orf64)). P. sansomeana* mitochondria also included a unique ORF predicted to encode a protein of 402 aa *(orf402)* (Fig. 1). No detectable similarity was found between *orf402* and any known genes in GenBank. All protein coding genes and ORFs had the TAA stop codon except the *nad11* gene, which had the TGA stop codon.

**Fig. 1.**
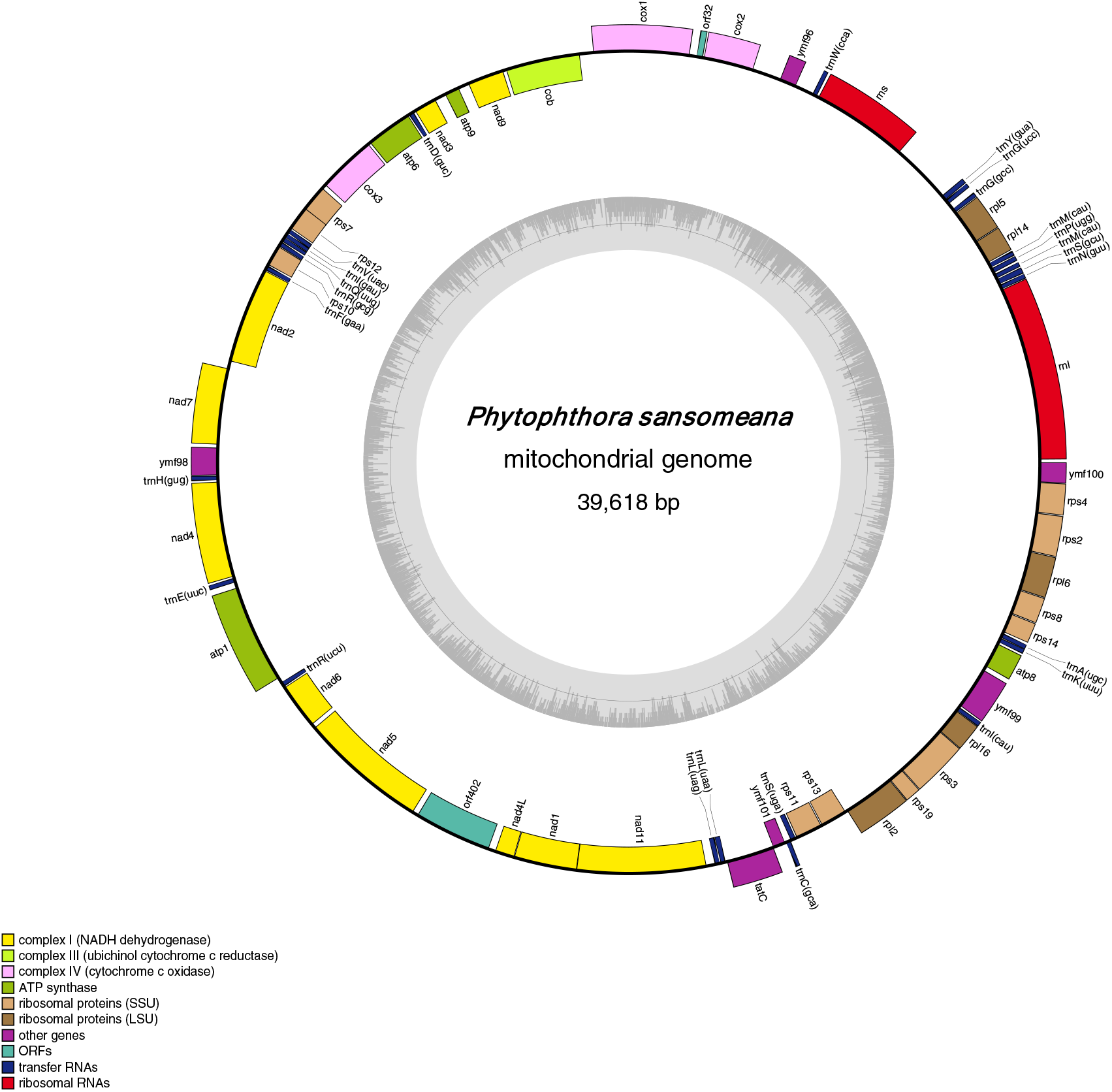
Circular map of the mitochondrial genome of *Phytophthora sansomeana.* Outer ring shows the predicted genes which are encoded on both strands. Inner ring shows local GC Density.

There was one large subunit rRNA gene *(rnl)* and one small subunit rRNA gene (rns). The 25 tRNA genes encoded tRNAs for 19 essential amino acids. There were two tRNA genes for arginine, glycine, leucine, methionine and serine, and one each for the other 14 essential amino acids except threonine (Fig. 1). There was a third tRNA gene located between *rpl16* and *ymf99* (Fig. 1) that had a CAU anticodon, but we interpreted it as *trnI_CAU_* rather than *trnM_CAU_* based on its similarity to a homologous tRNA gene at the same location in *P. infestans.* Experimentally confirmed in *Escherichia coli* [36] and inferred in *P. infestans* [14], it was assumed that the cytosine in *trnI_CAU_* anticodon was post-transcriptionally modified to lysidine that would enable it to recognize the AUA codon for isoleucine.

The genome was compact. Coding regions approximately comprised 90.1% of the genome. There was no intron in any of the genes. Of the 69 intergenic regions, 53 were 50 bp or less in length and 41 were less than 20 bp. There were three overlaps: between *rps7* and *rps12* (26 bp), between *nad1* and *nad11* (4bp), and between *tatC* and *ymf101* (4bp).

### Protein coding gene contents in the mitochondrial genomes of 10 *Phytophthora* species

Mitochondrial genomes of nine other *Phytophthora* species were publicly available and they were downloaded from GenBank (Table 1). Throughout the analysis below, please refer to Table 1 for reference and GenBank accessions. These 10 genomes are similar in size, ranging from 37,561 bp in *P. nicotianae* to 42,977 bp in *P. sojae.* G+C contents range from 21.57% in *P. polonica* to 22.39% in *P. ipomoeae.* Based on seven nuclear loci and four mitochondrial loci, *Phytophthora* species were grouped into 10 clades [23, 37]. The 10 species included in our analysis belong to clades 1, 7, 8 and 9 (Table 1).

All 10 genomes share the 34 genes with known functions, including the 14 genes in respiratory complexes, four ATP synthetase genes and 16 ribosomal RNA protein genes as described in *P. sansomeana.* The gene *atp9* was not reported in *P. nicotianae.* Our analysis of the submitted sequence in GenBank unambiguously identified *atp9* gene in this genome at the expected location (Fig. 2) but its 3’ end was truncated. DNA sequence alignment (not shown) revealed that this was caused by a deletion from the 3’ end of *atp9* gene that extended into the intergenic region between *atp9* gene and *nad9* gene. This deletion removed the last 13 amino acids in atp9 protein and replaced it with five non-homologous amino acids (Fig. 3).

**Figure 2.**
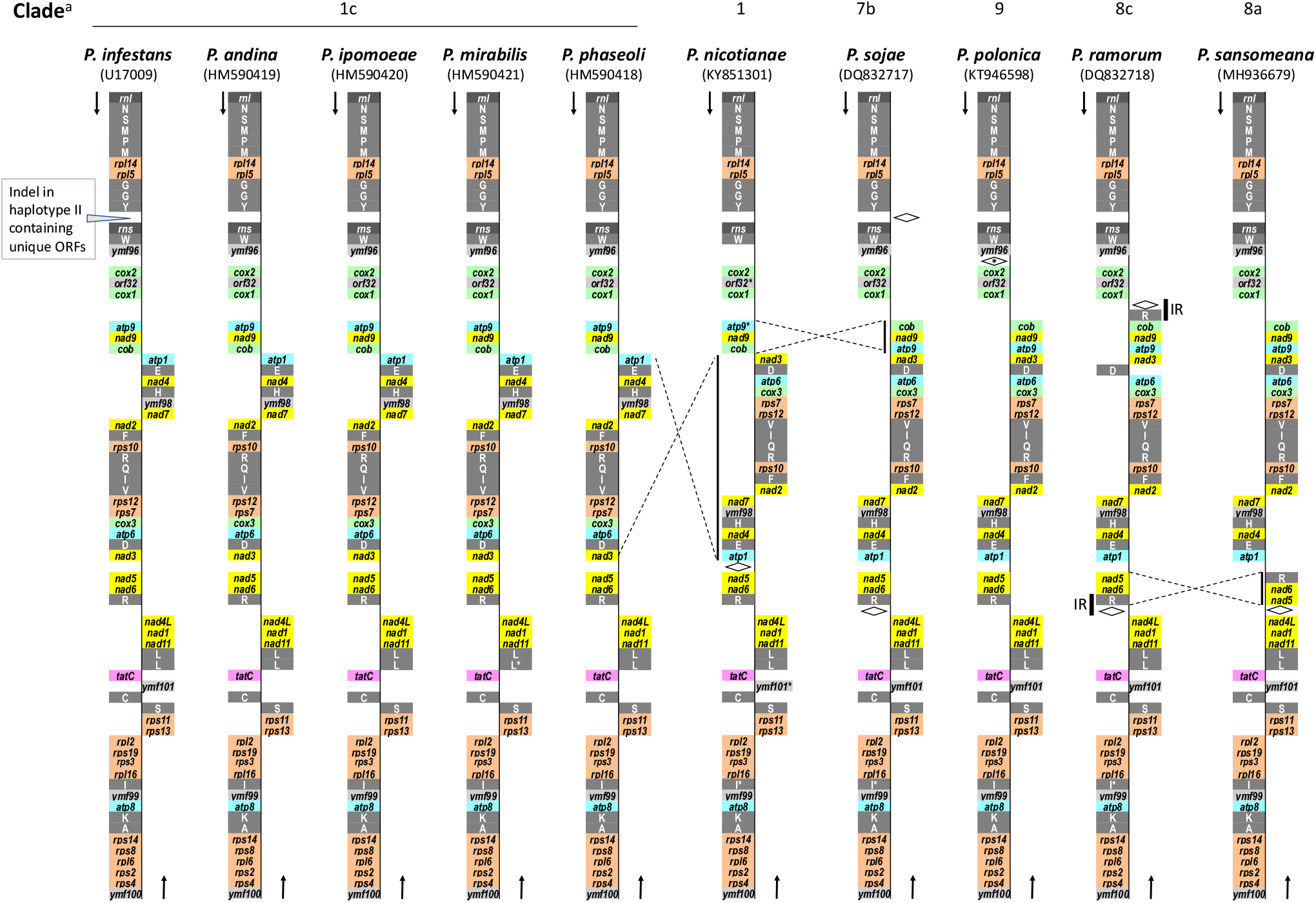
Linear representation of gene contents and organization of mitochondrial genomes of 10 *Phytophthora* species. RNA genes are in white letters with rRNA genes in dark grey background and tRNA genes in grey background. Protein coding genes are in dark letters with the following backgrounds: cytochrome C reductase and oxidases, green; ATP synthetases, blue; NADH dehydrogenases, yellow; ribosomal protein genes, brown; twin-arginine translocase, purple; and conserved ORFs with unknown function, light grey. The diamond symbols represent regions with one or more ORFs unique to that particular species. In species other than *P. sansomeana,* protein coding genes with an asterisk are those identified in this study not presented in GenBank annotations. tRNA genes with an asterisk are those whose annotations are modified in this study, either in coding strand or identity, comparing to GenBank annotations. IR, inverted repeat. ^a^ Clade designation is based on [23, 37].

**Fig. 3.**
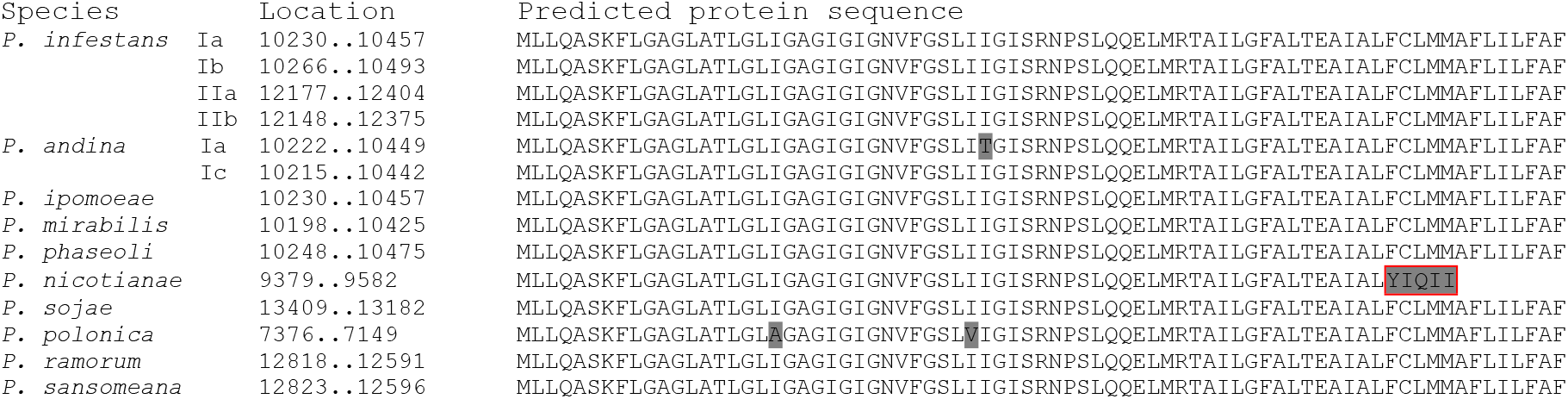
Deduced protein sequences of *atp9* gene in the mitochondrial genomes of 10 *Phytophthora* species. Amino acids differ from the consensus are shaded in grey. A deletion at the 3’ end of this gene in *P. nicotianae* results in the replacement of the last 13 amino acids with five non-homologous ones (boxed in red).

In addition to the 34 genes described above, another gene shared by these mitochondrial genomes was annotated with function, but its annotation in these genomes was inconsistent. *ymf16* (*=orf244)* is a conserved ORF in all 10 *Phytophthora* species that encodes a protein of approximately 244 amino acids. In the 13 mitochondrial genomes of nine *Phytophthora* species available in GenBank (Table 1), this gene was given the name *ymf16* in most cases and its product was annotated as SecY-independent transporter protein. However, in *P. ramorum* and *P sojae,* this gene was named *SecY,* and in *P. nicotianae,* its product was named SecY. The name “SecY” is not compatible with the meaning of “SecY-independent”.

There are three translocation systems in bacteria that transport distinct subset of proteins into or cross the inner membrane. SecYEG is the main translocase that transports unfolded peptides [38, 39], of which SecY is the main transmembrane subunit. YidC functions as an insertase and also assists SecYEG in the assembly of membrane proteins [40]. The twin-arginine translocation (Tat) system is capable of translocating folded or even multi-subunit protein complexes across inner membrane [41]. Proteins translocated by the Tat system have a twin-arginine motif in their signal peptides. The TatC protein is an essential component of the Tat system.

In a previous study, *TatC* gene was identified in the mitochondrial genomes of oomycetes [42]. To determine the function of *ymf16* gene, its predicted protein sequences from the 10 *Phytophthora* species were used to search Pfam database version 32 [43]. The only hit was TatC (sec-independent translocase protein, E values < 1e-20). As such, we annotated this gene as *tatC* (Fig. 2).

Six conserved ORFs have been reported in the mitochondrial genomes of these 10 species. Of these, *orf32, ymf98, ymf99* and *ymf100* were found in all 10 species. *Orf32* was not reported in *P. nicotianae* but our analysis found this ORF in the expected location (nucleotide 7530..7628) (Fig. 2). *Ymf96* was not found in *P. nicotianae.* Of the five species in subclade 1c, *ymf101,* a small ORF encoding approximately 64 aa, was found in *P. infestans* but missing in three species, *P. ipomoeae, P. mirabilis* and *P. phaseoli* (Fig. 2). Interestingly, In *P. andina, ymf101* was missing in haplotype Ic but our analysis identified it in haplotype Ia (nucleotide 30270..30476). *ymf101* was not reported in *P. nicotianae,* but our analysis identified it at nucleotide 30206..30015.

Unique ORFs have been reported in several species. In *P. infestans* haplotypes IIa and IIb, an insertion of approximately 2 kb encodes multiple ORFs [15]. Also reported were orf183 in *P. nicotianae,* two inverted copies of *orf176* in *P. ramorum,* and six ORFs at two location in *P. sojae.* Additionally, we identified *orf402* in *P. sansomeana* and also found an ORF with 173 codons (excluding the stop codon) in *P. polonica* at nucleotide 540..1061 (Fig. 2).

### RNA gene contents in the mitochondrial genomes of 10 *Phytophthora* species

Each genome had two rRNA genes for the large and small subunit ribosomal RNAs *(rnl* and *rns,* respectively). All the genomes encoded 25 tRNA genes and these genes had the same anticodons (Table S1). The only exception was *P. ramorum,* in which an inverted repeat of 1, 150bp resulted in a second copy of *trnR_UCU_* and *orf176* (Fig. 2 and Table S1). None of the mitochondrial genomes encoded tRNA for threonine. A tRNA gene for threonine was not found in the mitochondrial genomes of other oomycetes including *Phythium insidiosum* [35], *Phythium ultimum* [34], *Peronospora effuse* [44], *Pseudoperonospora humuli* [45], *Pseudoperonospora cubensis* [46], *Saprolegnia ferax* [47], *Achlya hypogyna* and *Thraustotheca clavate* [48]. Two stramenopiles, *Thalassiosira pseudonana* [49] and *Phaeodactylum tricornutum* [50], also lacked tRNA genes for threonine in their mitochondrial genomes, suggesting that this is a shared trait of mitochondria in stramenopiles.

### Organization of the mitochondrial genomes of 10 *Phytophthora* species

As discussed above, the mitochondrial genomes in these 10 species were conserved in gene content with few exceptions. These genomes were also conserved in gene order and coding strand except for three inversions. One inversion event occurred within clade 1. *P. nicotianae* belongs to clade 1 and is basal to subclades 1b and 1c. Compared to *P. nicotianae* and species in clades 7, 8 and 9, a large section between *atp1* gene and *nad3* gene was inverted in the five species in subclade 1c: P. *andina, P. infestans, P. ipomoeae, P. mirabilis* and *P. phaseoli.* This section contained 11 protein coding genes and eight tRNA genes. A second inversion occurred between species in clade 1 and species in clades 7, 8 and 9. This inversion covered three protein coding genes: *atp9, nad9* and *cob.* A third inversion occurred within clade 8. In *P. sansomeana* (subclade 8a), the section covering *nad5, nad6* and *trnR_UCU_* was inverted comparing to *P. ramorum* (subclade 8c) and species in other clades (Fig. 2).

### Phylogeny of *Phytophthora* species based on mitochondrial genomes

Other then *atp9,* 34 protein coding genes with annotated functions and the two rRNA genes, *rnl* and *rns,* were used to construct the phylogenetic tree (Fig. 4). Previously, *Phytophthora* species were grouped into 10 clades [23, 37]. Our analysis included species from four clades and the result was in agreement with previous studies. Species from individual clades were grouped together with bootstrap support 86% or higher. Within clade 1, previous studies showed moderate support that *P. nicotianae* was basal to subclades 1b and 1c based on seven nuclear genes [37] and four mitochondrial genes [23]. Our analysis supported the conclusion that *P. nicotianae* was basal to subclade 1c (100% bootstrap support). Species in subclade 1b was not included in this study. Within clade 1c, *P. phaseoli* diverged first and was basal to other species. *P. andina* hyplotype Ia was grouped with hyplotype I of *P. infestans,* while hyplotype Ic was grouped with *P. ipomoeae* and *P. mirabilis.* These findings were in agreement with previous reports [16, 51].

**Fig. 4.**
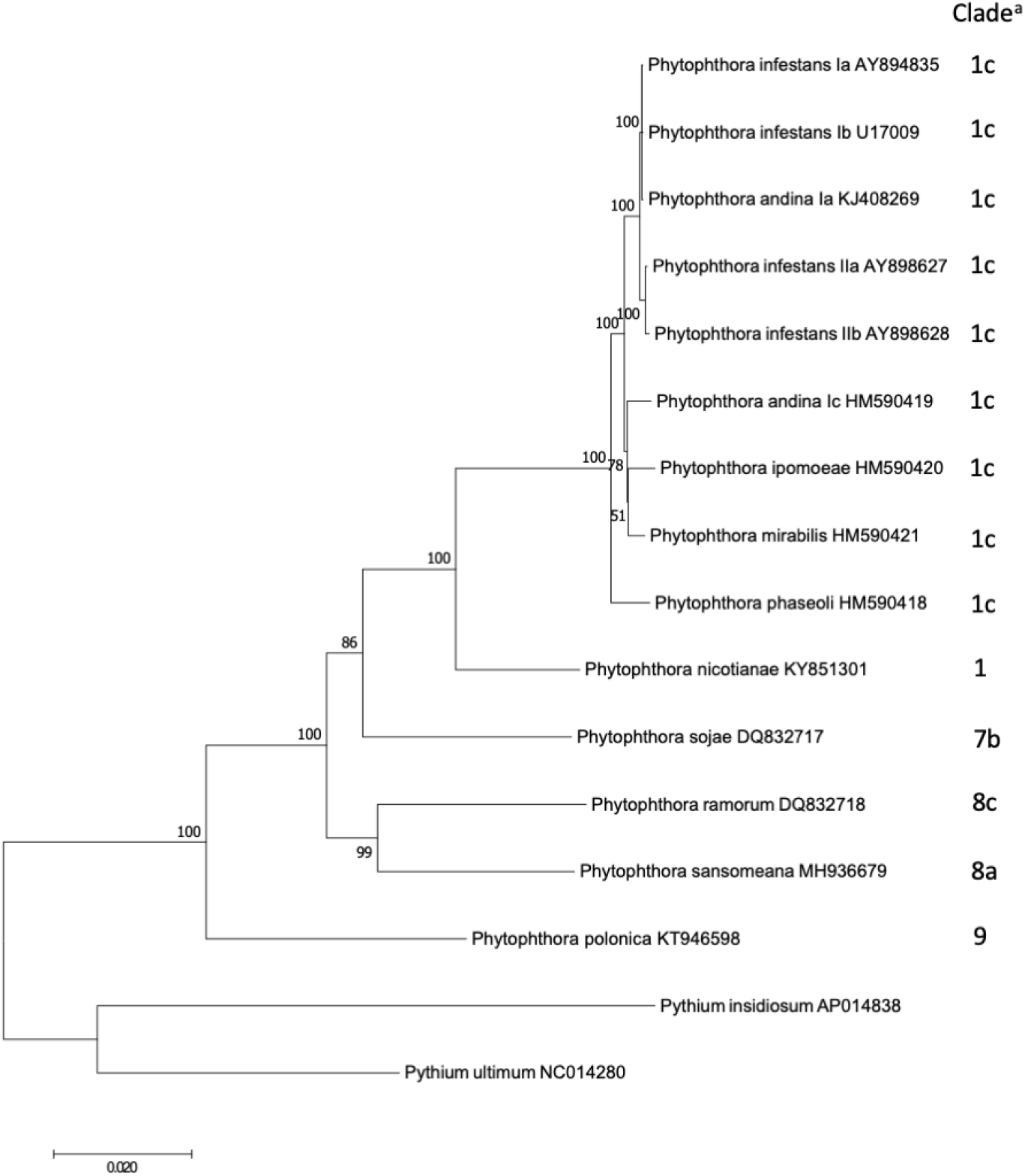
A maximum-likelihood tree based on the mitochondrial genomes. Branch support over 50% out of 1,000 bootstraps are shown. ^a^ Clade designation is based on [23, 37].

In summary, we sequenced and annotated the complete mitochondrial genome of the soybean pathogen *P. sansomeana* and compared it to that of nine other *Phytophthora* species. Inconsistencies in annotation among these mitochondria genomes were corrected. These genomes were found to be conserved in gene content with few exceptions. Three inversion events, each covering multiple genes, were observed among these genomes. Phylogenomic analysis using the mitochondrial genomes supported or enhanced previous findings.

Mention of trade names or commercial products in this publication is solely for the purpose of providing specific information and does not imply recommendation or endorsement by the U.S. Department of Agriculture. USDA is an equal opportunity provider and employer.

